# CRISPR/Cas9 gene drives in genetically variable and non-randomly mating wild populations

**DOI:** 10.1101/071670

**Authors:** Douglas W. Drury, Dylan J. Siniard, Gabriel E. Zentner, Michael J. Wade

**Affiliations:** Department of Biology, Indiana University, Bloomington, IN, USA 47405

## Abstract

Synthetic gene drives based on CRISPR/Cas9 have the potential to control, alter or suppress populations of crop pests and disease vectors, but it is unclear how they will function in wild populations. Using genetic data from four populations of the flour beetle *Tribolium castaneum*, we show that most populations harbor genetic variants in Cas9 target sites, some of which would render them immune to drive (ITD). We show that even a rare ITD allele can reduce or eliminate the efficacy of CRISPR/Cas9-based synthetic gene. This effect is equivalent to and accentuated by mild inbreeding, which is a characteristic of many diseasevectoring arthropods. We conclude that designing such a drive will require characterization of genetic variability and the mating system within and among targeted populations.

## Results and discussion

Gene drives based on engineered nucleases (hybrid meganucleases, ZFNs, TALENS and CRISPR/Cas9) offer a potentially significant improvement on sterile-male release methods for suppressing populations of agricultural pests and disease vectoring insects (*1-3*), allowing more rapid transformation of populations with fewer releases of modified insects (*4*). The RNA-guided CRISPR/Cas9 system permits the construction of custom synthetic gene drives potentially capable of spreading agriculturally valuable and medically beneficial genomic alterations made in laboratory organisms through wild populations (*5*). While initial studies of CRISPR-based gene drives in *Drosophila melanogaster* (*6*), *Anopheles stephensi* (*7*), and *Anopheles gambiae* (*8*) have demonstrated efficient drive propagation in lab-reared inbred populations, it is not clear how such gene drives will work in genetically diverse wild populations, where features like population structure, epistasis, and competition affect their evolutionary dynamics.

Gene drives work by segregation distortion (*9*) or ‘super Mendelian’ inheritance, wherein heterozygous individuals either transmit a desired gene in >90% of their gametes instead of the 50% Mendelian expectation or are transformed into homozygotes. Natural examples include *segregation distorter* in *Drosophila melanogaster (10)*, *Medea* in *Tribolium castaneum (11)*, the *t-haplotype* in *Mus musculus (12)*, and *spore killer* in *Neurospora crassa (13)*. In each natural case, the biased production of gametes of one allele is associated with some loss of fitness, so that some heterozygotes persist, which has allowed the drives to be discovered and studied. Indeed, a viability fitness loss of only 25% (selection coefficient, s, of - 0.25) in both sexes or a fertility loss of 50% in one sex is sufficient to counter even the most extreme transmission bias, preventing the spread of a driven allele (*14*). The upper limits to fitness loss are twice as high in germline gene-drive models (*15*) where individuals heterozygous at fertilization are converted to homozygotes. Although all models to date assume random mating, a small level of inbreeding significantly slows the rate of spread by diminishing the frequency of heterozygotes (see below). Because *any* gene in the genome that suppresses drive is favored by natural selection, Charlesworth and Charlesworth (*14*) concluded that ‘*a newly evolved distorter system will be inefficient*.’ Does this concern apply to laboratory-engineered CRISPR/Cas9 gene drives proposed for release into wild populations for purposes of population suppression? Here, we use evolutionary genetic analysis incorporating standing genetic variation and inbreeding in the model insect and agricultural pest *Tribolium castaneum* to address this question.

For the purpose of assessing the effects of naturally occurring genetic variation on CRISPR/Cas9-based drive efficiency, we consider three separate regions of Cas9 target sites: the protospacer-adjacent motif (PAM), the seed region, and the outer protospacer. The PAM (often NGG, recognized by the commonly used *Streptococcus pyogenes* Cas9), is required for Cas9 to recognize and cleave its target site (*16*). Thus, conversion of either G to another base in the NGG PAM would be severely deleterious to drive copying and we therefore call these variants in the PAM **Immune-To-Drive** variants or **ITD**s. The five bases immediately adjacent to the PAM, which are collectively known as the seed region (*17*), are responsible for nucleation of the gRNA:DNA hybrid, necessary for stable Cas9 binding and subsequent cleavage. Indeed, genome-wide characterization of Cas9 binding by ChIP-seq has shown binding but little cleavage at targets with seed region mutations (*17, 18*) as well as substantially decreased Cas9 binding in some cases (*18, 19*). Seed mutations are thus expected to severely compromise drive copy and for our modeling purposes we treat them ITDs. Third, in the protospacer outside of the seed region, Cas9 exhibits a relatively relaxed specificity (*20*). Thus, SNPs in this region would be expected to contribute minimally to drive resistance.

To assess the potential for standing genetic variation to contribute to drive resistance, we obtained the genomic sequence of high quality gRNA-targeting regions from each of four *T. castaneum* populations, Bhopal (India), Jerez (Spain), Purdue (Indiana), and Lima (Peru), for exons in three potentially drive-relevant genes. The first, *w*, encoding an ABC transmembrane transporter involved in eye pigmentation in *D. melanogaster* (*21*) was chosen as it has been used as a landing site for an anti-malarial gene drive in *A. stephensi (7)* and its disruption imparts a moderate fitness cost in *D. melanogaster* (*22*) and *A. stephensi (7)*. We found two SNPs in the selected *w* gRNA, one in the outer protospacer, present at low frequency in the Jerez population but moderate frequency in the Purdue population and one in the seed region, present at low frequency in the Purdue population (Figure 1). Due to the sequence flexibility of the outer protospacer, the first SNP would not be expected to substantially contribute to impaired Cas9 recognition and cleavage. However, the second SNP, in the seed region, could severely reduce Cas9 association and subsequent cleavage and so is considered an ITD. We next considered a gRNA in *Ace2*, involved in development, female fertility, and insecticide resistance in *T. castaneum* (*23*) and thus expected to carry a heavy fitness penalty when disrupted. We found a single SNP in the *Ace2* gRNA, present at low frequency in Purdue and moderate frequency in Peru. Importantly, this SNP lies at a critical base, the second G of the PAM (Figure 1), and so would be expected to completely abrogate Cas9 target site recognition and cleavage. However, it has recently been reported that NGA is the most effective non-canonical PAM for cleavage by *S. pyogenes* Cas9 (*24*), so this variant may not totally impair recognition and cleavage. However, because Cas9 cleavage even with an NGG PAM is not necessarily 100% effective and the NGA PAM is less robust in cell-based CRISPR assays, we treat this NGG-to-NGA substitution as a PAM-disabling ITD. The last gRNA targeted the *T. castaneum* homolog of the highly conserved (*25*) Sperm flagellar protein 1-encoding gene *TC010993*, mutation of which would be expected to severely reduce male fertility. The *TC010993* gRNA harbored a rare SNP present only in the Peru population in the outer protospacer region that would likely have little or no impact on drive spread (Figure 1). Although there are no ITDs in any of the three gRNA-targeting regions in the India population, other work (*26*) has shown that this population is reproductively incompatible with populations from North and South America.

**Figure 1.**
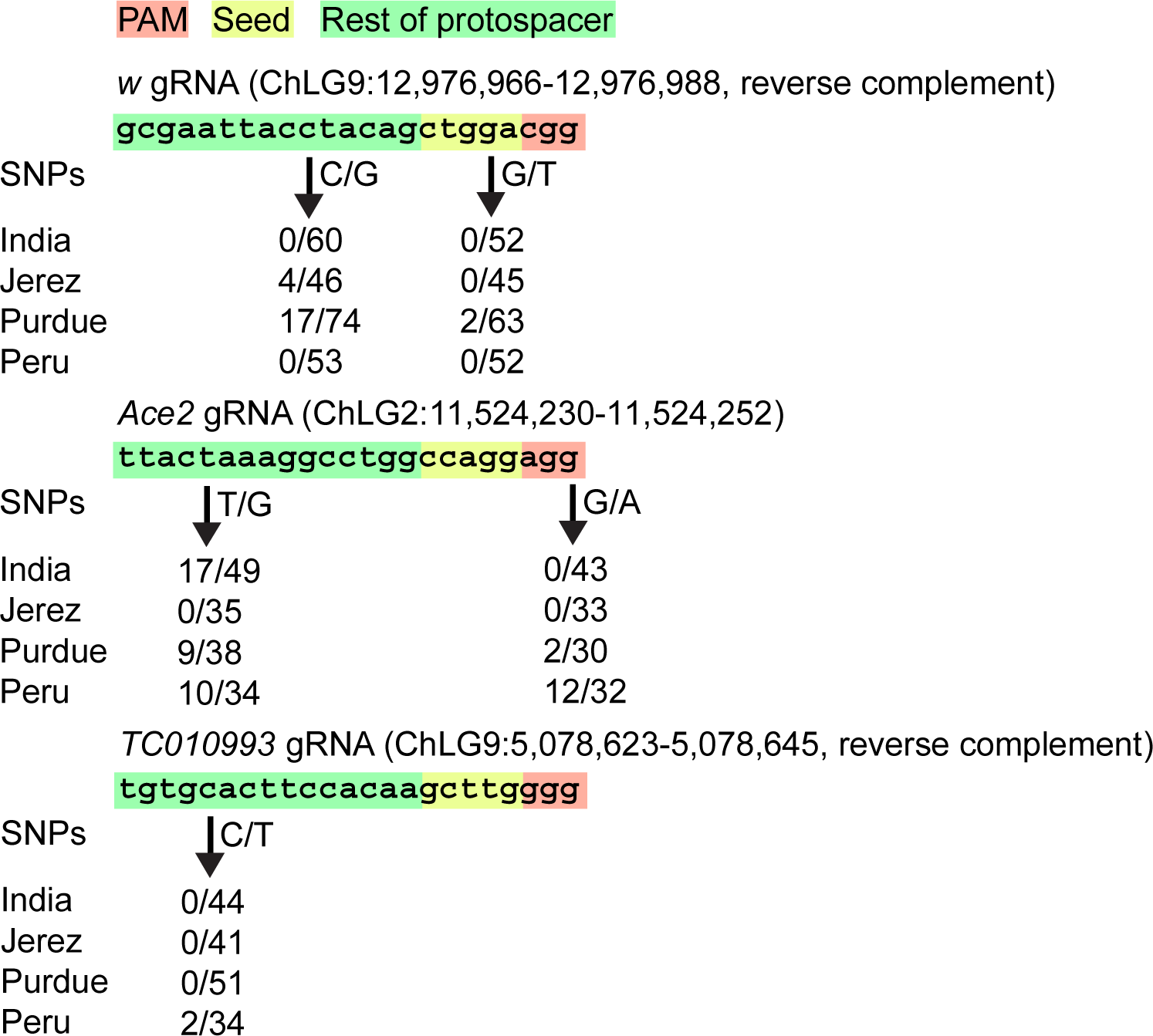
Genetic variation in Cas9 target sequences. Each component of the Cas9 target sequence is highlighted according to its potential to impact Cas9 cleavage (PAM = red, complete abrogation of cleavage; seed = yellow, reduced cleavage efficiency; outer protospacer = green, little to no effect on cleavage efficiency). The frequency of each SNP in the analyzed population is given below each gRNA sequence. Note that all regions harbor SNP variants in some population and that an obstructing variant in the critical PAM region of *Ace2* occurs at a frequency of 0.375 in the Peru population.

To see how these SNP variants affect gene drive and population suppression, we modified the recursion formula for q, the CRISPR-like drive-gene frequency, given in eq. (1) of (*15*) to allow for inbreeding and ITDs. The wild-type allele frequency is p and that of the ITD variant is r, where (q + p + r) = 1. In eq. (1) below, wild-type and ITD homozygous fitness is 1; individuals homozygous for the drive construct have fitness (1 - s); and, ITD/drive and wild-type/drive heterozygotes have fitness (1 - hs), where h is the degree of dominance. The rate at which wild-type alleles are converted to drive alleles in heterozygotes equals c, where 0 ≤ c, s ≤ 1. All individuals converted from heterozygotes to homozygotes have fitness of (1 – s). We let F be the proportion of individuals mating, non-randomly, within their own type; for example, 2q(1-q)(1-F) is the frequency of heterozygotes weighted by the proportion of individuals mating non-randomly. Half of the heterozygotes lost by inbreeding are added to the frequency of each respective homozygote; e.g., q^2^ becomes q^2^ + q(1-q)F. The frequency of the drive allele in the next generation is now 

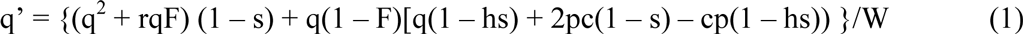
where mean fitness, W, equals 1 – s{q^2^ + rqF + 2rqh(1 – F) + 2pqc(1 – F) + 2pqh(1 – F)(1 – c)}. When F = r = 0, q’ and W reduce to the formulas of (*15*). Mean fitness of the population after the introduction of gene drive also measures the degree of population suppression. We thus modeled propagation of a CRISPR/Cas9-based drive based on each of the gRNAs described above. We considered each drive to be highly efficient (c = 0.95) and near dominant (h = 0.98). The *TC010993* drive is an exemplar of an ideal situation, in which a highly deleterious gene drive (s = 0.8) is introduced at a frequency of 0.7 into a randomly mating wild population harboring no ITDs. In this case, the drive is fixed after 8 generations, effectively suppressing the population (Figure 2A). However, when an ITD is present at low frequency (0.01) in a randomly mating population, similar to a drive scenario using the *Ace2* gRNA described above, the frequency of the drive rapidly declines while ITD frequency increases, until the drive frequency is < 0.01 and the ITD frequency is ~0.45 after 10 generations (Figure 2B). We note that the ITD frequency of 0.01 is lower than the frequency of the *Ace2* gRNA ITD in either of the populations in which it is present, suggesting that our estimate is conservative in this case. Lastly, we considered a scenario in which a gene expected to have lower fitness consequences is disrupted, perhaps with an anti-pathogen drive as has been described in *A. stephensi (7)*, which would be analogous to a drive based on the *w* gRNA we selected. We modeled this drive using all the above-described parameters except that s = 0.4 to represent a modest fitness effect. In this example, the drive is present at a frequency > 0.9 for 63 generations but declines to < 0.01 after 91 generations (Figure 2C), indicating that a low-frequency ITD can eventually halt the spread of a drive with a moderate organismal fitness penalty.

**Figure 2.**
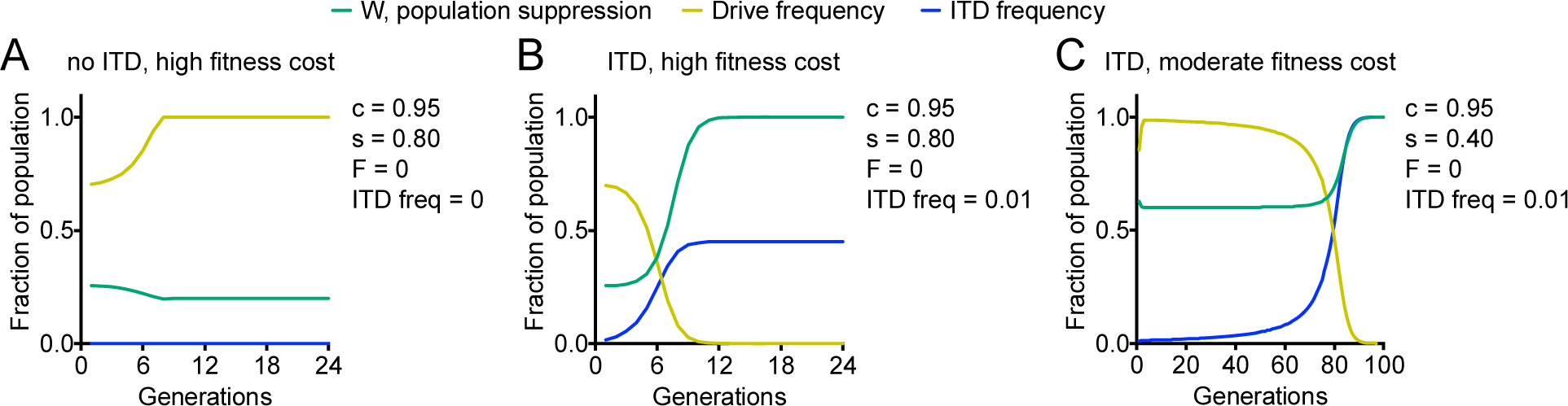
ITDs limit drive propagation. (A) In the absence of an ITD and inbreeding, a strongly deleterious drive rapidly spreads to fixation. (B) Addition of an ITD at a frequency of 0.01 severely impairs propagation of a strongly deleterious drive and leads to its removal from the population, as well as a substantial increase in ITD frequency. (C) In the presence of an ITD at a frequency of 0.01 a moderately deleterious drive remains at high frequency for several tens of generations but is eventually eliminated from the population.

We next tested the effect of inbreeding on drive propagation. While rates of inbreeding (F) in *T. castaneum* have not been determined, such measurements have been made for mosquitoes and range from not significant to 0.8 (*27-29*). For our model, we chose a conservative estimate of F, 0.15. In the absence of ITDs, as in the *TC010993*-like scenario described above, this degree of inbreeding resulted in a rapid crash of drive frequency, leading to a frequency of < 0.01 in 6 generations (Figure 3A). Combination of an ITD at a frequency of 0.01, similar to our *Ace2* example, with inbreeding at a rate of 0.15 had a negligible effect on the speed at which the drive crash occurred (Figure 3B). Notably, inbreeding also limited the spread of the ITD relative to the corresponding no-inbreeding scenario (Figure 2B), likely due to the rapidity of the cessation of drive spread (Figure 3B). Lastly, introduction of inbreeding into the *w*-like situation described in Figure 2C) led to a drastic acceleration of drive loss, with drive frequency reaching < 0.01 in 39 rather than 91 generations (Figure 3C).

**Figure 3.**
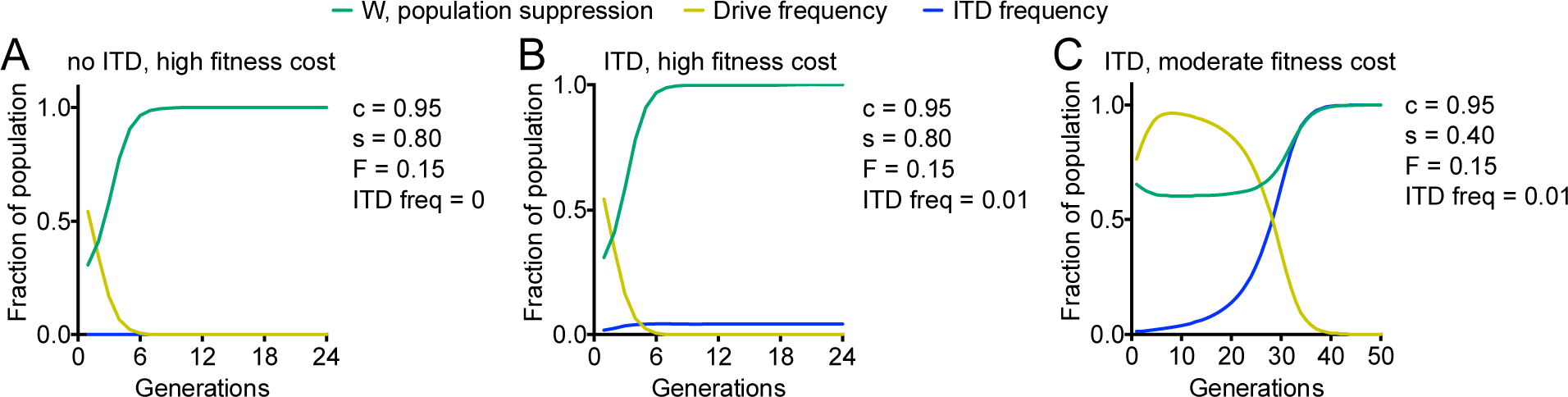
Inbreeding impairs drive spread. (A) Inbreeding at a rate of 0.15 in the absence of an ITD leads to rapid cessation the spread of a strongly deleterious drive and its removal from the population. (B) Addition of an ITD at a frequency of 0.01 to the population inbreeding at a rate of 0.15 again leads to rapid loss of the strongly deleterious drive from the population, as well as a moderate increase in ITD frequency. (C) In the presence of an ITD at a frequency of 0.01 a moderately deleterious drive remains high for several generations but is effectively eliminated (frequency <0.01) after 39 generations.

We have modeled the effects of standing genetic variation and inbreeding on CRISPR/Cas9-based gene drives using *T. castaneum* as a model. Our results suggest that even a low-frequency ITD can severely limit the spread of a highly deleterious drive and cause its elimination from a population. Moreover, such ITDs were selected for and were present at a higher frequency in the population following drive elimination, rendering the population more refractory to repeated releases of individuals carrying the same drive. We found that ITDs have a less pronounced effect on drives with a lower fitness cost to the organism but are still capable of eliminating the drive from a population over time. This is particularly relevant to anti-pathogen drives inserted into loci thought to have little fitness effect on an organism, such as the insertion of an anti-malarial drive into the *white* locus of *A. stephensi* (*6*), emphasizing the importance of minimizing fitness effects on the host. We found that, in the case of a strongly deleterious gene drive, moderate inbreeding caused a very rapid loss of the drive regardless of the presence of an ITD. Furthermore, Bull (*30*) has shown that gene drive can favor a gene that increases the tendency to inbreed, so the use of a constant level of inbreeding in our modeling is conservative. Inbreeding also accelerated the loss of a moderately deleterious drive, but the drive was stable for several generations before declining.

In sum, our analyses suggest that standing genetic variation and inbreeding could have wide-ranging and sometimes severe consequences on the efficiency of drive propagation in a wild population. Inbreeding, which in our analyses had the most profound impact on drive propagation, may be most effectively overcome by saturation of the target population with drive individuals through repeated, high-volume releases. To minimize the potential impacts of ITDs on drive spread, propose that a survey of genetic diversity at the locus of interest in the target population is a necessary first step in designing a CRISPR-based gene drive. If a target region of a locus of interest is highly variable, it may be prudent to select a different region of the gene to be disrupted to avoid the potential complications of ITDs described here. However, if targeting of a region with an altered PAM is highly desirable, a Cas9 variant engineered to alter its PAM specificity may be used (*31*). While the impediments posed to CRISPR/Cas9-based gene drives described here are substantial, we are confident that with appropriate consideration of population structure and genetic diversity such drives can be effectively used to spread beneficial genetic modifications through wild populations.

## Materials and methods

### gRNA design

Exon sequences for *w*, *Ace2*, and *TC010993* (Tcas3 assembly) were obtained from BeetleBase (http://beetlebase.org) (*32*) and input into CRISPRdirect (https://crispr.dbcls.jp) (*33*) using the Tcas3 assembly for the specificity check. Only high-quality gRNA sequences (defined as having a single 20mer + PAM and a single 12mer + PAM match) were selected for SNP analysis.

### Populations and Sample Preparation

A total of 96 individuals were randomly collected from four wild-caught laboratory populations of *T. castaneum* (Bhopal, India; Lima, Peru; Lafayette, IN (Purdue); Jerez, Spain). Each population was established from individuals collected from local populations of *T. castaneum* found in granaries and markets. All populations were collected and are maintained at population sizes of greater than 250 individuals to reduce the potential for inbreeding and loss of genetic variation in the laboratory. To compare the genetic variation within and between populations, DNA was extracted and purified using the CTAB method from 24 randomly selected individuals from each population (*34*). Each individual DNA sample was uniquely barcoded using Nextera Indexing Kit and pooled for sequencing. Three individual DNA samples were discarded due to low yield after library preparation and fragment size selection steps.

### Sequencing and mapping

We applied 150-bp paired-end Illumina (Illumina HiSeq platform) sequencing to 93 individuals (24 individuals each from India, Peru, Purdue; 21 individuals from Jerez). Each individual was sequenced to a coverage depth of ~2-3X with an average of ~50X total coverage for each population. The Fastq files were filtered using Cutadapt (https://cutadapt.readthedocs.io/en/stable/) to remove low quality sequence and the resulting files were processed for mapping using Fix Pairings (https://github.com/svmzhang/make_life_easier). The paired-end reads for each individual were to against the Tcas3 reference genome using BWA 0.7.6a (*35*). The resulting SAM files were converted to BAM and mpileup files using SAMtools (*36*). mpileup files were synced using PoPoolation2 (https://sourceforge.net/projects/popoolation2/). From this, we obtained a direct major and minor allele count for every SNP in our target loci.

## Acknowledgements

This work was supported by Indiana University startup funds (G.E.Z.) and National Institutes of Health R01 GM084238 (M.J.W.).

